# Rapid inactivation of SARS-CoV-2 with Deep-UV LED irradiation

**DOI:** 10.1101/2020.06.06.138149

**Authors:** Hiroko Inagaki, Akatsuki Saito, Hironobu Sugiyama, Tamaki Okabayashi, Shouichi Fujimoto

## Abstract

The spread of novel coronavirus disease 2019 (COVID-19) infections worldwide has raised concerns about the prevention and control of SARS-CoV-2. Devices that rapidly inactivate viruses can reduce the chance of infection through aerosols and contact transmission. This *in vitro* study demonstrated that irradiation with a deep ultraviolet light-emitting diode (DUV-LED) of 280 ±5 nm wavelength rapidly inactivates SARS-CoV-2 obtained from a COVID-19 patient. Development of devices equipped with DUV-LED is expected to prevent virus invasion through the air and after touching contaminated objects.

## Letter

The novel coronavirus SARS-CoV-2 pandemic has spread worldwide and placed countries in emerging, rapidly transforming situations. The World Health Organization (WHO) clarified that more than 5.3 million cases of COVID-19 and 342,000 deaths had been reported to WHO by 25 May 2020 [1]. Infectious virus is detected in specimens from the respiratory tract, nasopharyngeal sites, and feces in COVD-19 patients [2]. Recently, infectious SARS-CoV-2 was isolated from the urine of a COVID-19 patient [3]. SARS-CoV-2 is detectable in aerosols for up to 3 h, up to 4 h on copper, up to 24 h on cardboard and up to 2–3 days on plastic and stainless steel [4]. To prevent exposure to contaminated material (contact infection), which is one of the major transmission routes, hand hygiene with alcohol is recommended, but its effectiveness in preventing the spread of SARS-CoV-2 infection may be insufficient [5, 6].

A deep ultraviolet light-emitting diode (DUV-LED) instrument generating around 250–300 nm wavelength has been reported to effectively inactivate microorganisms, including bacteria, viruses and fungi [7–10], but effects on SARS-CoV-2 have not been reported. We evaluated the antiviral efficacy of irradiation by DUV-LED, generating the narrow-range wavelength (280±5 nm) (Nikkiso Co., Tokyo, Japan), against SARS-CoV-2.

A strain of SARS-CoV-2 isolated from a patient who developed COVID-19 in the cruise ship *Diamond Princess* in Japan in February 2020 [11] was obtained from the Kanagawa Prefectural Institute of Public Health (SARS-CoV-2/Hu/DP/Kng/19-027, LC528233). The virus was propagated in Vero cells cultured in minimum essential medium (MEM) containing 2% fetal bovine serum (FBS). At 48 h after infection, virus stocks were collected by centrifuging the culture supernatants of infected Vero cells at 3,000 rpm for 10 min. Clarified supernatants were kept at -80 °C until use. Aliquots of stock virus were diluted with phosphate-buffered saline and adjusted to 2.0 ×10^4^ plaque-forming units (PFU)/ml. For the evaluation of DUV-LED inactivation, aliquots of virus stock (150 μl) were placed in the center of a 60-mm Petri dish and irradiated with 3.75 mW/cm^2^ at work distance 20 mm for a range of times (n=3 each for 1, 10, 20, 30, or 60 s). Each virus stock irradiated with DUV-LED was serially diluted in 10-fold steps, then inoculated onto Vero monolayers in a 12-well plate. After adsorption of virus for 2 h, cells were overlaid with MEM containing 1% carboxymethyl cellulose and 2% FBS (final concentration). Cells were incubated for 72 h in a CO_2_ incubator, then cytopathic effects were observed under a microscope. An unirradiated virus suspension was used as a negative control. To calculate PFU, cells were fixed with 10% formalin for 30 min, followed by staining with 0.1% methylene blue solution. The antiviral effects of DUV-LED irradiations were assessed using the logPFU ratio, calculated as logPFU ratio=log_10_ (Nt/N0), where Nt is the PFU count of the UV-irradiated sample, and N0 is the PFU count of the sample without UV irradiation. In addition, the infectious titer reduction rate was calculated as (1-1/10^log^ ^PFU^ ^ratio^) × 100 (%). All experiments were performed in a BSL-3 laboratory.

We observed a marked cytopathic effect in virus-infected cells without DUV-LED irradiation (Figure 1A, see “0 s”). In contrast, virus-infected cells irradiated for 60 s showed largely comparable morphology to mock cells (Figure 1A, see “60 s”). To our surprise, virus-infected cells irradiated for 1 s showed minimal change (Figure 1A, see “1 s”). The plaque assay (Figure 1B) revealed that short time DUV-LED irradiation rapidly inactivated SARS-CoV-2 (Figure 1C and Table S1). Of note, the infectious titer reduction rate of 87.4% was already recognized with irradiation of virus stock for 1 s, and the rate was 99.9% with irradiation for 10 s. These results suggest that DUV-LED drastically inactivated SARS-CoV-2 with irradiation for even a very short time.

**Figure 1.**
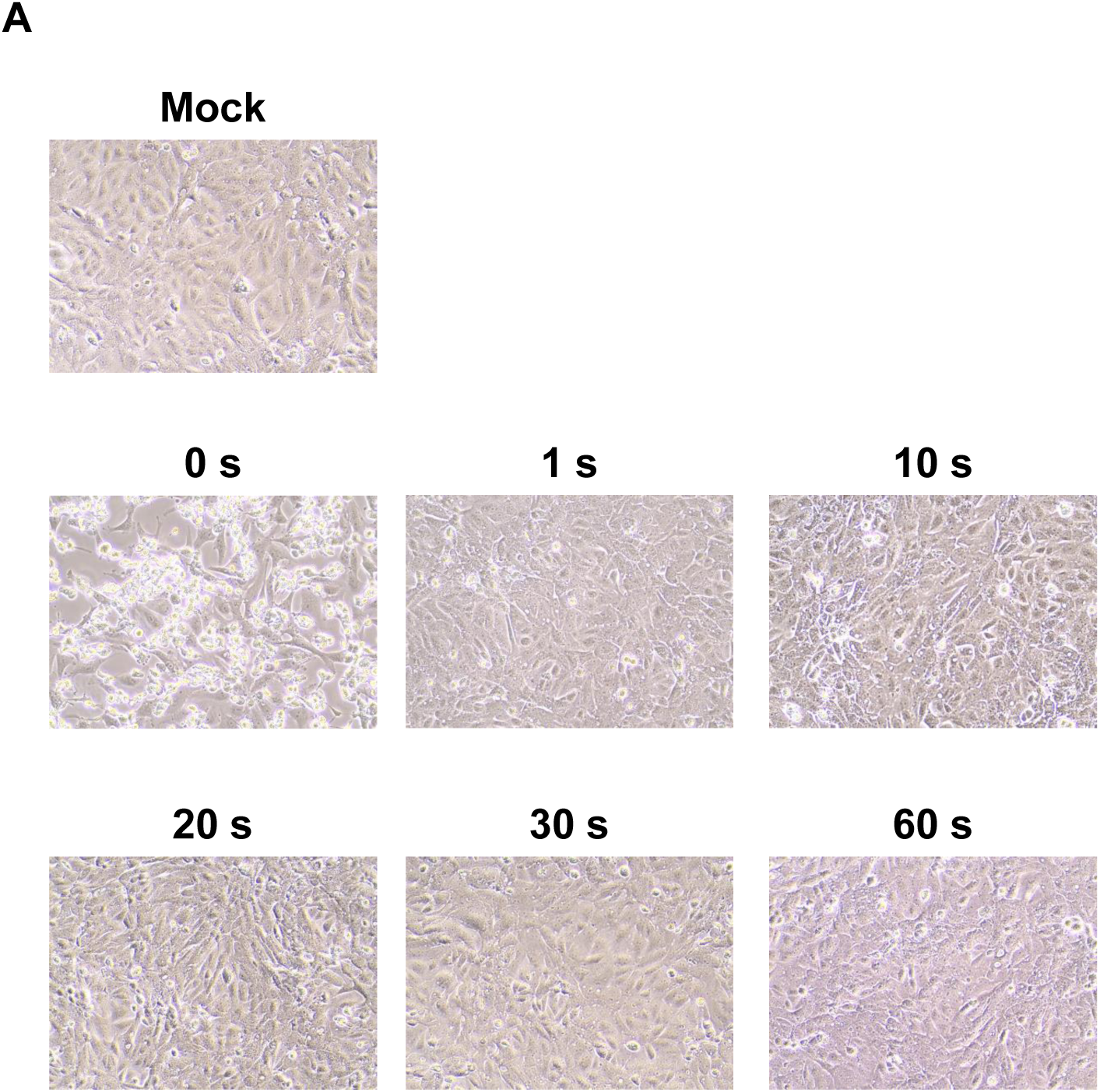
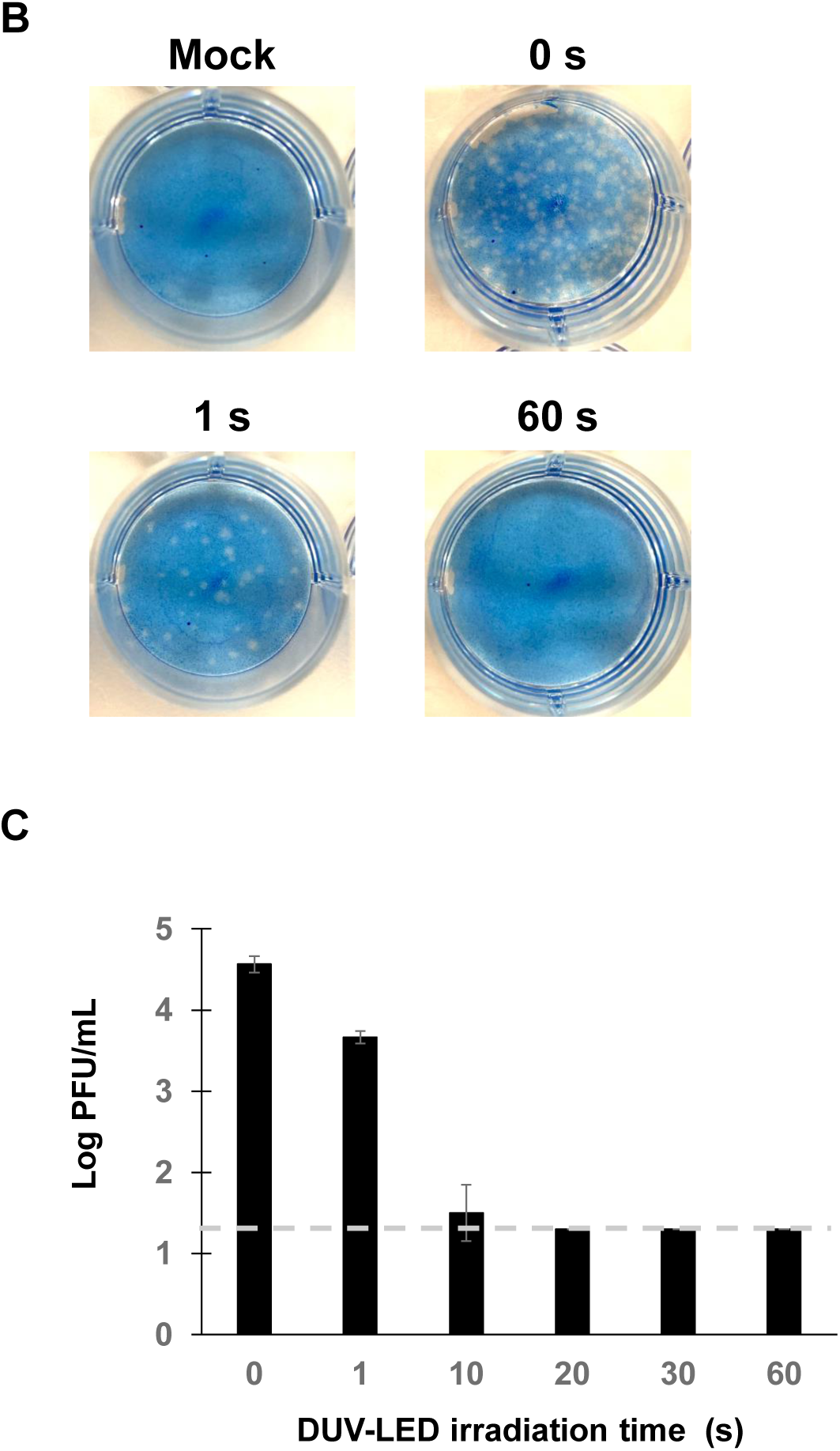
Inhibitory effects of DUV-irradiation on SARS-CoV-2. (A) Cytopathic changes in virus-infected Vero cells without DUV-LED irradiation (0 s), or with DUV-LED irradiation for 1, 10, 20, 30 or 60 s. (B) Plaque formation in Vero cells. Virus solutions irradiated with DUV-LED for several durations were diluted (100-fold) and inoculated to Vero cells. A representative result is shown. (C)Time-dependent inactivation of SARS-CoV-2 by DUV-LED irradiation. The results shown are the mean and standard deviation (SD) of triplicate measurements. The dashed line indicates the limit of detection.

UV-LEDs providing irradiation at various peak emission wavelengths, such as UV-A (320–400 nm), UV-B (280–320 nm), and UV-C (100–280 nm), have been adopted to inactivate various pathogenic species, including bacteria, viruses and fungi. Devices equipped with UV-LEDs are now beginning to be introduced into medical fields. UV-C is considered to be the most effective germicidal region of the UV spectrum, acting through the formation of photoproducts in DNA [12]. These pyrimidine dimers interrupt transcription, translation and replication of DNA, eventually leading to inactivation of microorganisms [13]. The efficacy of this inactivation may depend not only on the wavelength, but also on factors such as the target (e.g., bacterial species), light output and environmental conditions. The DUV-LED we used has the characteristics of a narrow-range wavelength and high power for short exposure times and long-term use. This study demonstrated for the first time the rapid inactivation of SARS-CoV-2 under DUV-LED irradiation. As shown in Figure 1B, cytopathic effects were observed in control Vero cells infected with SARS-Cov-2, but not in these cells with DUV-LED irradiation for only 10 s. As well as in community settings, healthcare settings are also vulnerable to the invasion and spread of SARS-CoV-2, and the stability of SARS-CoV-2 in aerosols and on surfaces [4] likely contributes to virus transmission in medical environments. No vaccines, neutralizing antibodies, or drugs are currently available for prevention and treatment of SARS-CoV-2. By revealing that SARS-Cov-2 inactivation can be achieved with very short-term DUV-LED irradiation, this study provides useful baseline data toward securing a safer medical environment. Development of devices equipped with DUV-LED is expected to prevent the virus invasion through the air and after touching contaminated objects.

## Contributors

H.I. and H.S. conceived the study and wrote the manuscript. A.S. and T.O. conducted the experiments dealing with viruses. S.F. contributed to the study design, study supervision and manuscript revision.

## Acknowledgements

We wish to thank Drs. Tomohiko Takasaki and Jun-Ichi Sakuragi, from the Kanagawa Prefectural Institute of Public Health, for providing the SARS-CoV-2/Hu/DP/Kng/19-027 strain.

---

**Table S1.** Differences in infectious titer with different DUV-LED irradiation times.

| Irradiation time | control<br>(no irradiation) | DUV-LED irradiation time |  |  |  |  |
| --- | --- | --- | --- | --- | --- | --- |
|  |  | 1 sec | 10 sec | 20 sec | 30 sec | 60 sec |
| PFU(PFU/mL) | $3.7 \times 10^4$ | $4.7 \times 10^3$ | $2.7 \times 10^1$ | $6.7 \times 10^0$ | <20 | <20 |
| Log PFU ratio <sup>1)</sup> | — | 0.9 | 3.1 | >3.3 | >3.3 | >3.3 |
| Infection titer reduction ratio<sup>2)</sup>(%) | — | 87.4 | 99.9 | >99.9 | >99.9 | >99.9 |

<sup>1)</sup> $\log_{10} (N_t/N_0)$ , where $N_t$ is the PFU count of the UV-irradiated sample, and $N_0$ is the PFU count of the sample without UV irradiation. <sup>2)</sup> $(1 - 1/10^{\log \text{PFU ratio}}) \times 100$ (%).

